# Continuous Measures of Decision-Difficulty Captured Remotely: II. Webcam eye-tracking reveals early decision processing

**DOI:** 10.1101/2023.06.06.543799

**Authors:** Jennifer K. Bertrand, Alexandra A. Ouellette Zuk, Craig S. Chapman

## Abstract

As decisions require the gathering of relevant information, eye-tracking measures that capture the way visual information is typically acquired offer powerful indices of the dynamic decision-making process. This study is the second of a pair of studies that explore continuous measures of decision-making using remote, online tools in naturalistic settings. While cursor-tracking, used in the companion paper (Ouellette Zuk et al., 2023), enabled access to dynamic decision processes expressed during movement, in the present study, we now employ webcam eye-tracking to examine the dynamics of information gathering during decision making prior to movement initiation. Using three previously published binary choice tasks, we explored indices of decision difficulty in the gaze dynamics that would complement the motor measures in our companion paper. We find that harder choices elicit more eye dwells and longer final dwells, reflecting a decision resolution process that Ouellette Zuk et al. index during the final choice movement. Beyond this, we identify distinct gaze patterns uniquely employed in each task, revealing the utility and sensitivity of gaze metrics in illuminating the early difficulty-independent information gathering processes at play. Together, this paper series demonstrates the power of remote, online methods as tools for deeply understanding the complete, dynamic and continuous decision process, from the first glance to the final response.

## 1 Introduction

In the past, research on decision-making has primarily focused on the outcomes of decisions and has used discrete measures such as choice-outcomes, accuracy, or reaction time to understand the underlying cognitive processes involved (see [39] for review). However, these approaches really only capture *what* decision was made with little or no information about *how* a particular decision was arrived at. To address this gap, recent research has turned to the motor system as a means to investigate the dynamics of decision-making. Actions that move through space have been shown to track decision processes that unfold across time [7, 15, 11, 48].

One of the most accessible ways to record movement data is through cursor movements on a screen. For example, 2-D computer-mouse movements provide a sensitive, flexible, and scalable method to study decision dynamics ([26, 33, 21, 14, 12, 46], and many more). Moreover, the widespread adoption of cursor-based technologies (computers, tablets, smartphones) and the recent availability of online experiment generation tools (e.g., Labvanced [13], Gorilla [2], lab.js [22], jsPsych [32], PsychoPy [36]) means that now, more than ever, we can collect information about decision dynamics from larger, more diverse samples and from people in more ecologically-valid contexts (i.e., remote, online data collection). The current study is the second half of a two-article set aimed at mapping the possibilities and limitations of using remote data collection during cursor-based decision tasks. In the first study (Ouellette Zuk et al., 2023), we focus on testing the robustness of remote data collection for understanding dynamic decision-making, showing that nuanced details of decision processes are available not only in computer-mouse movements but are also clearly evident and sometimes even stronger when the same task is deployed on tablets and smartphones.

But, our first study is literally only half the story. While cursor-tracking reveals valuable insights into the decision process once movement begins, it cannot track details of the perceptual processes that occur prior to a movement toward a particular choice being initiated. While reaction time reliably tracks decision difficulty in this early phase (e.g., [38, 35]) it fails to capture any of the constituent dynamics of how the decision process is evolving. In these crucial early moments of a decision, a person is gathering the necessary information from their environment to inform their choice. For tasks involving visual stimuli in particular, this information gathering is primarily mediated by eye movements. Therefore, eye-tracking offers a unique opportunity to investigate the decision process at an earlier stage, revealing how individuals extract information and make decisions based on where they focus their gaze.

Given this, it is not surprising that eye-tracking has long been a prominent method in decision-making research. Extensive work has shown the interconnectedness of gaze and choice, where gaze patterns can both reflect and bias choices [42, 17, 16]. Moreover, eye movements have been shown to actively sample the world in a way that adaptively maximizes the informative value of fixations [19, 6]. Analyzing these fixation sequences that precede a decision has provided valuable insights, revealing that the location, duration, and pattern of fixations serve as indices of the relative competition between choice options [28, 29]. Findings like these have challenged classic decision-making theories (e.g., evidence accumulation; [18, 37, 43]) to reconcile the important role the eyes play in information sampling. This has led to the development of gaze-aware decision models such as the attentional drift diffusion model (aDDM; [28]), Decision Field Theory [5], and the gaze cascade model [42]. These models assume an option receives more evidence when gazed at, acting like an amplifier for the attended option [27]. Moreover, the aDDM has been extended beyond preferential choice contexts, encompassing different stimuli types (e.g., numeric information vs pictorial images; [30]) and choice domains (e.g., risky and social choices; [44]). Collectively, these theoretical advancements and empirical contributions provide a foundation for comprehending how the eyes sample spatially-distributed information, emphasizing the value of eye-tracking as a method to better understand how decisions unfold. In the context of the current study, they show that eye-tracking is almost a perfect complement to cursor-tracking in its ability to fill in the gap of decision dynamics during the earliest stages of a choice being made.

A major drawback shared by all of the aforementioned eye-tracking studies is their confinement to laboratory settings. Recently, however, the use of webcam eye-tracking has emerged as a promising avenue in bridging the gap between controlled laboratory experiments and data collected in a wide range of environments (e.g., [49, 47, 4]). Admittedly, webcam eye-tracking is still a method in its infancy and has notable limitations in both temporal and spatial accuracy [40, 3, 49]. Despite these challenges, it offers a distinct advantage by capturing gaze patterns in ecologically-valid settings such as within the participants’ own homes and on their personal devices. Furthermore, it can be argued that - with the overall trend toward increasing digital and screen based interactions - computerized decision-making tasks like the one used by Krajbich et al. [28], now closely resemble everyday, real-world decisions. Thus, by shifting these tasks to a more naturalistic context, such as the participants’ homes, a more comprehensive exploration of authentic, real-world visual behaviours and decision-making processes becomes possible. Additionally, the remote nature of this method allows for scalable data collection beyond the limitations of a laboratory, while also providing access to more diverse, global populations [23, 1].

Therefore, in this study we employ webcam eye-tracking - in part to explore its potential and limits as a method - and theoretically to explore the relationship between decision difficulty and gaze behaviour patterns. As previously mentioned, the current study is a companion paper to Ouellette Zuk et al. (2023), with both studies intentionally sharing the same experimental design. This design aimed to replicate and extend three unique and previously published mouse-tracking based decision-making tasks. These tasks were deliberately chosen to cover a range of decision domains including objective perceptual judgments (Numeric Size-Congruity [12]), semi-subjective conceptual judgements (Sentence Verification [10]) and subjective preference judgements (Photo Preference [26]). The tasks also varied in terms of stimulus characteristics, encompassing numerical digits, written statements, and photos. By employing webcam eye-tracking, we not only gain insights into how the decision context varies across tasks but also how differences in the presentation and distribution of information across space affect the decision-making process, something that cannot be solely obtained through mouse-tracking. Thus, replicating Ouellette Zuk’s design with eye-tracking will not only allow us to explore the rich, dynamic decision process earlier in time, beginning before movement initiation, but also explore how this process presents across decision contexts and with different distributions of decision information in the display. In doing so, we also demonstrate the utility of remote data collection in general and webcam eye-tracking in specific as a tool for capturing this rich readout of decision-making from participants in their own environments using their own devices.

## 2 Results

This study replicates and extends our cursor-tracking-focused companion paper (Ouellette Zuk et al., 2023). Both studies employed three binary choice tasks where participants indicated their choice through cursor movements: a Sentence Verification task [10], a Numeric-Size Congruity task [12], and a Photo Preference task [26]. Each task was designed and analyzed to produce Easy and Hard trials (see Figure 1): For the Sentence Verification task, based on previous work [10], participants indicated if a simple statement was true or false. On Easy trials, the statements were true and not negated (e.g., ‘Cars have tires’) and on Hard trials the statements were true and negated (e.g., ‘Cars do not have wings’). For the Numeric-Size Congruity task, based on previous work [12], participants indicated which of two digits had a higher numeric value. On Easy trials the size and numeric value were congruent (e.g., 2 vs. 8)), and on Hard trials size and value were incongruent (e.g., 2 vs. 8). Finally, for the Photo-Preference task, based on previous work [26], participants indicated which of two photos they preferred. On Easy trials, one photo had low pleasantness while the other had high pleasantness and on Hard trials both photos had high pleasantness. Our companion paper successfully replicated the main finding from the original task publications (i.e., [10, 12, 26]) that responses on Hard trials take longer than Easy trials, while also showing how decision difficulty affects cursor movements. Here we predict that we will also replicate the finding that Hard trials generate longer response times than Easy trials and investigate how decision difficulty affects webcam-tracked gaze behaviour.

**Figure 1:**
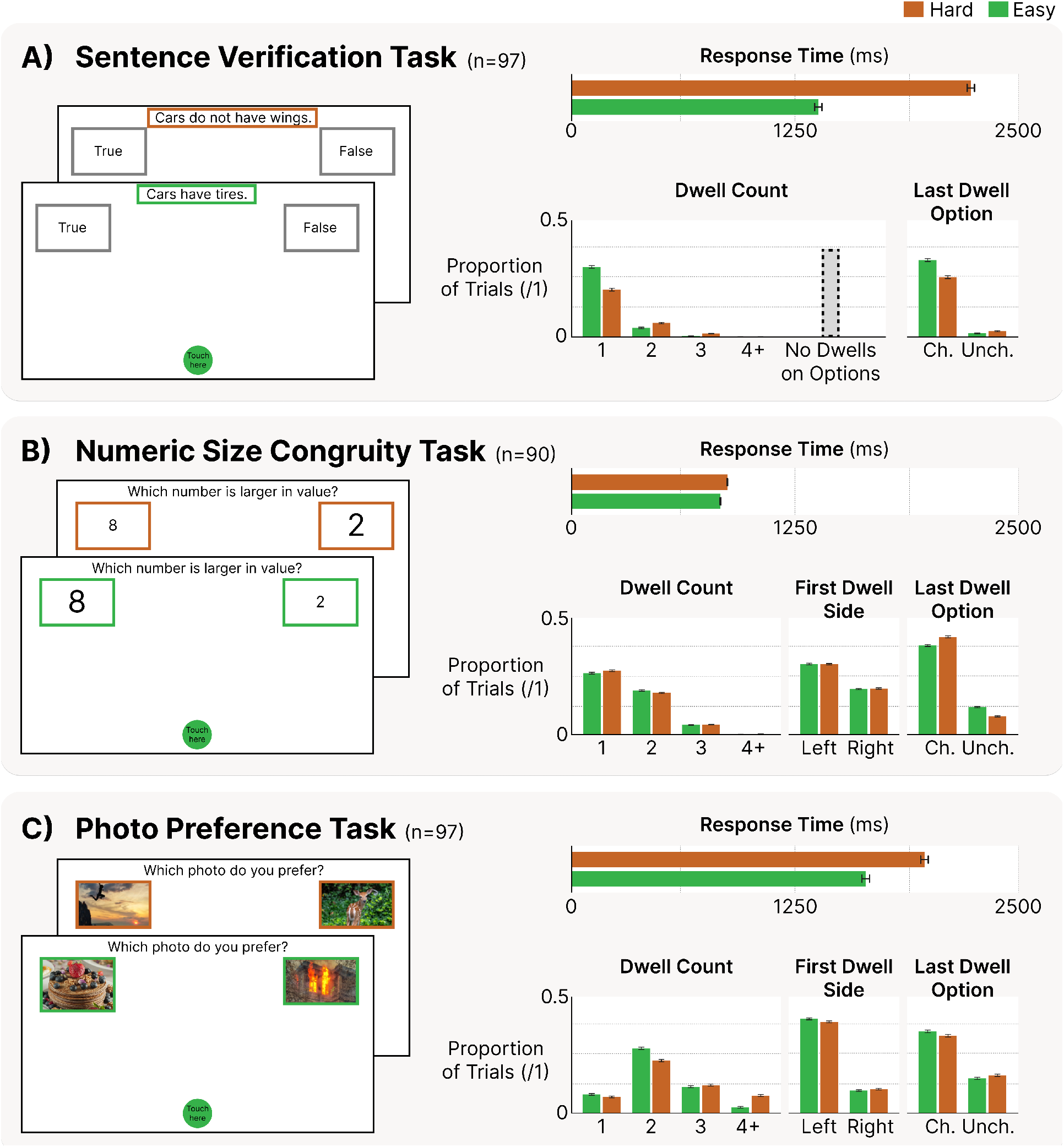
Examples of hard and easy decisions, alongside response time (horizontal bar graphs) and the proportion of trials (vertical bar graphs) results across the study’s three tasks: A) Sentence Verification, B) Numeric Size Congruity and C) Photo Preference. Throughout, orange represents hard decisions, while green represents easy decisions. Error bars are the standard error of the difference between the hard and easy conditions. The proportion of trials data is presented as simplified marginal means, where each factor analyzed in the Repeated Measures ANOVA (RMANOVA) is presented independently across its levels (note (Ch.) means Chosen and (Unch.) means Unchosen) The Sentence Verification task includes an untested but present gaze behaviour of No Dwells on Options, indicating the proportion of trials where only the sentence received a dwell.

### 2.1 Response Time: Hard decisions take longer than easy decisions

We use Response Time as a single, broad measure to capture the duration from the presentation of choice options to the point of response (marked by the cursor entering the selected option). It encompasses both reaction time and movement time as described in our companion paper (Ouellette Zuk et al., 2023). Replicating our previous work, and confirming our key prediction, across all three tasks we observed a consistent Difficulty effect (see Figure 1). Specifically, Hard trials required significantly more time than Easy trials (Sentence Verification: *t* (96) = 21.0, *p <* .001; Numeric-Size Congruity: *t* (89) = 8.01, *p <* .001; Photo Preference: *t* (96) = 7.95, *p <* .001).

### 2.2 Proportion of Trials: Unique task demands drive unique gaze behaviours

We began our examination of gaze patterns within each task by characterizing and analyzing the most frequently observed dwell patterns (see Figure 1). For our analysis, we defined a dwell as a continuous gaze on an area of interest, lasting at least 100 milliseconds within the expanded boundaries of that area (see Figure 3). A dwell ended if the gaze shifted outside that area for more than 100 milliseconds. To identify the most common gaze patterns we conducted separate RMANOVAs for each task. These analyses aimed to assess when and how often the eyes dwelled on particular AOIs. The following RMANOVAs include up to four factors to describe the dwell patterns observed: Dwell Count, Difficulty, First Dwell Side, and Last Dwell Option. The Dwell Count factor consisted of four levels (1, 2, 3, and 4 or more), indicating the number of unique dwells during a trial. The Difficulty factor distinguished between trials classified as either Hard or Easy. The First Dwell Side factor described whether the initial dwell on a trial fell upon the Left or the Right choice option. Finally, the Last Dwell Option factor indicated whether the final dwell of the trial was on the Chosen or Unchosen option.

**Figure 2:**
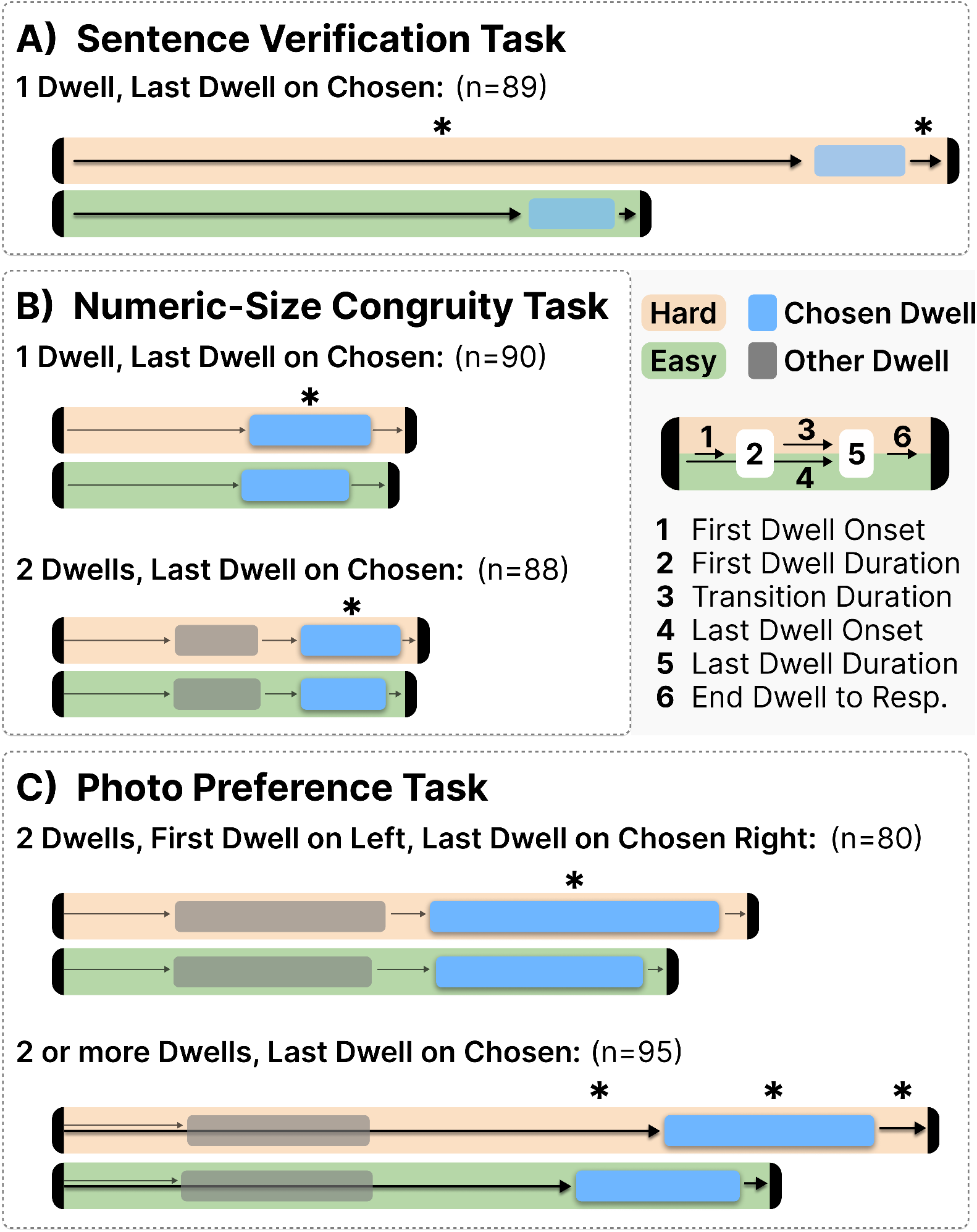
Gaze dynamics of the most common dwell patterns across the three decision tasks. Each pattern is depicted for hard and easy decisions, with every decision shown within a horizontal orange (hard) or green (easy) bar. All gaze patterns are aligned to the moment choice options are presented. Arrows are used to indicate the onsets and offsets of dwells, while the blue bar shows the dwell duration on the chosen option, and the grey bar shows the dwell duration on the other option for patterns where there is more than one dwell. Any significant differences between the gaze metrics of hard and easy trials are indicated with a star (*) and an opaque bar or thicker arrow. Insignificant differences are shown as transparent bars or thin arrows. For the Photo Preference 2 or more dwells plot, only the first and last dwell are depicted, but the results include trials where there would be additional dwells. Detailed means and statistics are available in Table 2.

**Figure 3:**
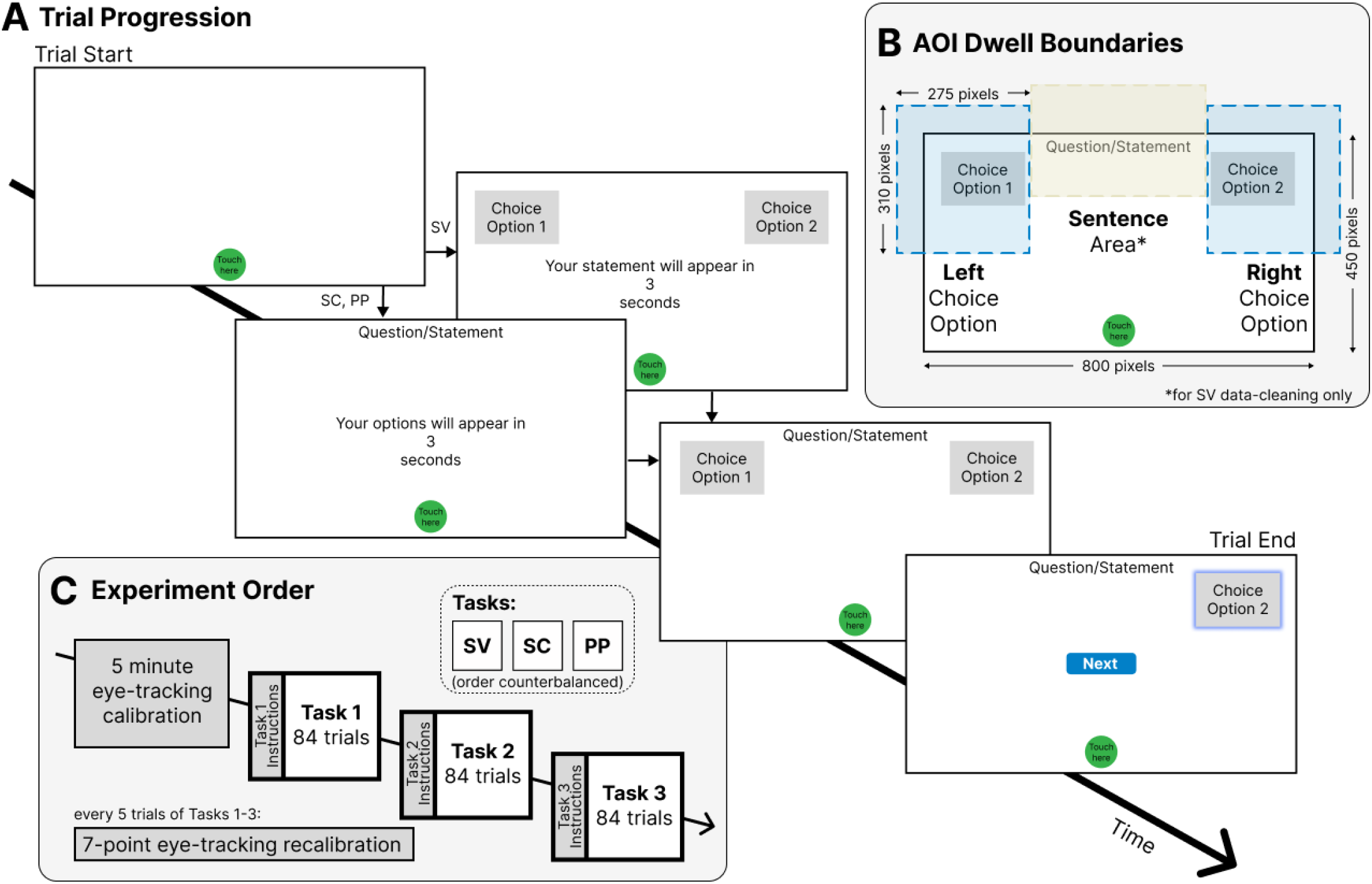
A) All three tasks presented a classic reach-decision paradigm requiring participants to choose one of two stimuli presented at the top left and right of their computer screen. For Numeric-Size Congruity (SC) and Photo Preference tasks (PP), countdown onset was accompanied by a question specific to the task, appearing at the top center of the display. The Sentence Verification (SV) task presented the two choice options coincident with countdown onset and presented a statement (rather than a question) upon countdown completion. Participants proceeded in a self-paced manner, pressing a button to begin the next trial. B) The enlarged areas of interest (AOIs) used to define the boundaries of the Left and Right choice options (blue transparent areas) when determining whether a dwell was made. The dimensions of the choice AOIs (275 × 310 pixels) are presented relative to the dimensions of the frame size (800 × 450 pixels). The Question/Statement AOI was used only for data-cleaning in the SV task. This frame was scaled to the size of each participant’s screen. C) Overview of the experiment’s design. Each participant completed an SV task, an SC task, and a PP task, with task order counterbalanced between participants. Eye-tracking calibration occurred at the beginning of the session, and a re-calibration procedure was performed every 5 trials. Task-specific instructions were presented prior to each task.

#### 2.2.1 Sentence Verification Task

Sentence Verification was unique in that many times (∼35% of all trials) participants’ eyes dwelled only on the sentence and never on either choice option. We represent the proportion of these “No Dwells on Options” in Figure 1A, but it is not possible to examine them in our statistical analysis of choice option gaze patterns. Thus, we acknowledge their presence as a predominant gaze behaviour in Sentence Verification and proceed with the rest of the analysis looking only at trials having at 1 or more dwells.

A 2 (Last Dwell Option) x 2 (Difficulty) x 4 (Dwell Count) RMANOVA of the proportion of trials within the Sentence Verification task revealed a significant three-way interaction between the tested factors (see Table 1; *F* (1.27,121.97) = 53.6, *p <* .001). This interaction was interrogated further by splitting the data into the much more common trials that ended with a dwell on the Chosen option (∼60%) and those that more rarely ended with a dwell on the Unchosen option (∼5%). For each of these groups we ran separate 2 (Difficulty) x 4 (Dwell Count) RMANOVAs.

**Table 1:** Proportion of trials results from each task’s three-way interaction involving Difficulty as a factor. Note. *p *<* .05; **p *<* .005; ***p *<* .0005

| Last Dwell<br>(%, SD) | Dwells<br>Count | Difficulty |  | Hard-Easy |  |  |  |  |
| --- | --- | --- | --- | --- | --- | --- | --- | --- |
|  |  | Hard | Easy | *M* | *SE* | df | *F* | *p* |
| Sentence Verification |  |  |  |  |  |  |  |  |
| Chosen<br>(58.78%, 27.53%) | 1 | 0.181 | 0.286 | -0.104 | 0.013 | 1,96 | 64.8 | *** |
|  | 2 | 0.057 | 0.038 | 0.019 | 0.006 | 1,96 | 10.9 | ** |
|  | 3 | 0.015 | 0.006 | 0.009 | 0.003 | 1,96 | 11.5 | ** |
|  | 4+ | 0.002 | 5.52e-4 | 0.002 | 9.83e-4 | 1,96 | 4.8 | * |
| Unchosen<br>(4.33%, 4.76%) | - | 0.006 | 0.004 | 0.002 | 0.001 | 1,96 | 5.2 | * |
| Numeric-Size Congruity |  |  |  |  |  |  |  |  |
| Chosen<br>(80.10%, 10.78%) | 1 | 0.230 | 0.195 | 0.035 | 0.008 | 1,89 | 20.1 | *** |
|  | 2 | 0.144 | 0.143 | 0.002 | 0.006 | 1,89 | 0.08 | n.s. |
|  | 3 | 0.042 | 0.042 | 4.32e-4 | 0.003 | 1,89 | 0.02 | n.s. |
|  | 4+ | 0.003 | 0.003 | 4.82e-4 | 9.79e-4 | 1,89 | 0.24 | n.s. |
| Unchosen<br>(19.90%, 10.78%) | 1 | 0.044 | 0.070 | -0.026 | 0.004 | 1,89 | 49.0 | *** |
|  | 2 | 0.034 | 0.048 | -0.013 | 0.003 | 1,89 | 16.5 | *** |
|  | 3 | 0.002 | 0.001 | 0.001 | 8.09e-4 | 1,89 | 2.7 | n.s. |
|  | 4+ | 0.00 | 0.00 | 0.00 | 0.00 | 1,89 | NaN | NaN |
| Photo Preference (only Left Start; 79.66%, 22.43%) |  |  |  |  |  |  |  |  |
| Chosen<br>(55.17%, 19.54%) | 1 | 0.032 | 0.029 | 0.003 | 0.005 | 1,96 | 0.44 | n.s. |
|  | 2 | 0.087 | 0.150 | -0.064 | 0.007 | 1,96 | 74.1 | *** |
|  | 3 | 0.091 | 0.088 | 0.003 | 0.007 | 1,96 | 0.16 | n.s. |
|  | 4+ | 0.053 | 0.022 | 0.031 | 0.006 | 1,96 | 27.9 | *** |
| Unchosen<br>(24.48%, 13.05%) | 1 | 0.008 | 0.017 | -0.009 | 0.003 | 1,96 | 8.6 | ** |
|  | 2 | 0.089 | 0.079 | 0.010 | 0.007 | 1,96 | 1.9 | n.s. |
|  | 3 | 0.016 | 0.017 | -0.001 | 0.004 | 1,96 | 0.13 | n.s. |
|  | 4+ | 0.016 | 0.002 | 0.014 | 0.003 | 1,96 | 18.5 | *** |

When we look at the three-way interaction follow-up RMANOVA for Last Dwell on Chosen we reveal a significant interaction between Difficulty and Dwell Count (*F* (1.29,123.74) = 56.4, *p <* .001), as well as significant main effects (Difficulty: *F* (1.33,127.76) = 281.9, *p <* .001; Dwell Count: *F* (1,96) = 41.1, *p <*.001). We further followed up the two way interaction with four 1-factor RMANOVAs - one for each level of Dwell Count. Here, we see significant Difficulty effects in the proportion of trials at each level of Dwell Count. For last dwell on Chosen one-dwell trials, we see a higher proportion of Easy trials than Hard trials (*p <* .001, *M* _*Hard−Easy*_ = -0.104), while the reverse pattern of a higher proportion of Hard than Easy trials is revealed when there are two, three, and four or more dwells (all *p*’s *<* .05, and *M* _*Hard−Easy*_ range from 0.00215 to 0.0192). In general, this suggests that for trials that show the commonly-occurring Last Dwell on Chosen pattern, Harder trials result in more dwells than Easy trials.

When the Last Dwell fell on the Unchosen option (which occurred rarely, *M* _*Unchosen*_ = 0.0434), we observed differences in the proportion of trials driven by both Difficulty (*F* (1,96) = 5.19, *p* = .0249) and the Dwell Count (*F* (1.17,112.37) = 61.16, *p <* .001), but not their interaction (*F* (1.22,116.65) = 1.99, *p* = 0.1582). The proportion of Hard trials was slightly higher than that of Easy trials (*M* _*Hard−Easy*_ = 0.0024), and all pairwise comparisons between the number of dwells yielded significant results (all *p*’s *<* .05), with the proportion of trials following a pattern of one-dwell *>* two-dwells *>* three-dwells *>* four or more dwells. The results suggest that participants have very few trials with multiple dwells in this task and that the relatively rare gaze pattern where the eyes end on the Unchosen option occurs slightly more on Hard trials than Easy trials.

Overall, Fig 1A highlights the general gaze patterns elicited by the Sentence Verification task: while sometimes there was no gaze upon the choice options, when it did happen, it was usually only once, and almost always on the chosen option. However, in the relatively infrequent number of trials where the gaze dwelled more than one time on the options, this occurred more often on hard trials.

#### 2.2.2 Numeric-Size Congruity Task

We explored the proportion of trials in Numeric-Size Congruity with a 2x2x2x4 RMANOVA (Last Dwell Option x First Dwell Side x Difficulty x Dwell Count). Two significant three-way interactions were revealed (these were the highest order significant interactions): Last Dwell Option x Difficulty x Dwell Count (see Table 1; *F* (1.85,164.29) = 22.2812, *p <* .001) and Last Dwell Option x First Dwell Side x Dwell Count (*F* (2.26,201.05) = 18.0261, *p <* .001).

To follow-up the Last Dwell Option x Difficulty x Dwell Count interaction, like the Sentence Verification follow-up, we looked at Difficulty x Dwell Count at each Last Dwell Option level separately. Again, participants were much more likely to end the trial by looking at the Chosen (∼80%) as compared to the Unchosen (∼20%) option. Following up the three-way interaction for the more common last dwell on Chosen gaze pattern, we uncovered a two-way interaction between Difficulty and Dwell Count (*F* (1.75,155.67) = 10.1,*p <* .001). The further follow-ups at each level of Dwell Count found that the interaction was driven primarily by a significant Difficulty effect on one-dwell trials only. That is, there is a greater proportion of single, Chosen option dwells on Hard trials compared to Easy trials (*p <* .001, *M* _*Hard−Easy*(1*Dwell*)_ = 0.0353). Overall, this result highlights the high proportion of trials where the last dwell ends on the chosen option, while revealing a subtle difficulty effect in single dwell trials. We believe this difficulty effect is best understood after also explaining the pattern of behaviour on Last Dwell on Unchosen trials.

For the less common Last Dwell on Unchosen trials, the 2x4 follow-up RMANOVA revealed a significant two-way interaction (*F* (1.86,165.17) = 26.9, *p <* .001). Further follow-ups highlight that this is an effect driven by a significantly greater proportion of Last Dwell on Unchosen trials in Easy one and two-dwell cases than Hard (both *p*’s *<* .001, *M* _*Hard−Easy*(1*Dwell*)_ = -0.0261, *M* _*Hard−Easy*(2*Dwells*)_ = -0.0133). Last Dwell on Unchosen is a less common gaze pattern as compared to Last Dwell on Chosen, but when present, it’s more likely to happen on Easy trials where there’s only one or two dwells to the choice options. So, why are we seeing fewer dwells on harder trials, opposite to what we saw in the other tasks? We speculate this has to do with a unique property of the Numeric-Size Congruity task where what makes a specific trial hard is the physical size of the target, which we feel is likely related to its visual discriminability. Specifically, on Hard trials, your eyes have landed on a physically small but numerically large number. This smaller character likely takes additional time to resolve. During this time, however, it is possible that you are also processing information from the other target location [45]. On these Hard trials, the other numeral at this peripheral location is numerically small and physically large. We think that it may be easier to resolve this peripheral, larger target, eliminating the need for a second fixation. If we take the mirror scenario, on an Easy trial your eyes land on a numerically large digit that is also physically large. Resolving this larger stimulus occurs quickly. But, the peripheral target in these cases is physically small. We speculate that on some trials there is too much uncertainty about the identify of the peripheral stimulus which in turn drives a second dwell to its location. The net result of this is that, on some Hard trials your linger at the first location and don’t need a second fixation while on some Easy trials you can more quickly leave the first location but feel the need to take a look at the second location.

Returning back to the second three-way interaction (Last Dwell Option x First Dwell Side x Dwell Count) that emerged from the omnibus RMANOVA, we chose to first explore this interaction at each level of First Dwell Side separately. In general, whether looking at trials where the gaze first fell on the Left option (which occurred ∼60% of the time) or the Right option (which occurred ∼40% of the time), the results are relatively similar. In both cases, a significant two-way interaction emerges between Last Dwell Option and Dwell Count (First Dwell Left: *F* (1.66,147.96) = 80.3, *p <* .001; First Dwell Right: *F* (1.74,154.92) = 149, *p <* .001). Further follow-ups for each case (at each level of Dwell Count) show the same pattern: the proportions of Last Dwell on Chosen trials is significantly greater than the Last Dwell on Unchosen trials (all 8 final tests have *p* values *<* .01).

All together, Figure 1B summarizes these results with marginal means shown for simplicity. In the Numeric-Size Congruity task, most often there were only one or two gazes upon the choice options, and in both cases, the first gaze was most likely to start on the left while the last gaze was almost always on the chosen option. The difficulty effects in this task were more subtle. When the gaze only landed on the chosen option and stayed there, it was more likely that this happened on a hard trial than an easy trial. But, in the small proportion of trials where the last dwell was on the unchosen option, there were more one and two dwell gazes on easy trials than hard trials. We speculate this difficulty effect has to do with peripheral processing and the visual discriminability of targets of different sizes.

#### 2.2.3 Photo Preference Task

We tested the proportion of trials in the Photo Preference task with a 2x2x2x4 RMANOVA (Last Dwell Option x First Dwell Side x Difficulty x Dwell Count) and found a significant four-way interaction between Last Dwell Option, First Dwell Side, Difficulty and Dwell Count (*F* (2.65,254.36) = 29.94, *p <* .001). For Photo Preference trials the most dominant factor was the First Dwell side with first looks to the Left (∼80%) being much more common than first looks to the Right (∼20%). As such, our initial follow-ups to the omnibus RMANOVA involved performing a 3-factor RMANOVA at each of the two levels of First Dwell Side.

In the more commonly-occurring trials where the First Dwell started on the Left, the three-way follow-up test revealed a significant three-way interaction between Last Dwell Option, Difficulty and Dwell Count (see Table 1; *F* (2.57,247.20) = 30.43, *p <* .001). The additional follow up two factor RMANOVAs that were performed for each level of Last Dwell Option revealed further interaction effects between Difficulty and Dwell Count in both tests (Last Dwell Unchosen: *F* (1.98,189.71) = 4.82, *p* = .009; Last Dwell Chosen: *F* (2.69,258.31) = 35.89, *p <* .001). In the most commonly occurring First Dwell Left-Last Dwell Chosen case, the two-way interaction follow-ups for each level of Dwell Count revealed a difficulty effect only for two and four or more dwells. For First Dwell Left, Last Dwell Chosen, two-dwell trials, there was a greater proportion of Easy trials than Hard trials (*M* _*Hard−Easy*_ = -0.0637, *p <* .001), and an opposite pattern for four or more dwell trials (*M* _*Hard−Easy*_ = 0.0312, *p <* .001). In the less common First Dwell Left, Last Dwell Unchosen case, when further follow-ups were performed at each level of Dwell Count, we found the two-way interaction to be driven by significant differences in trials with one and four or more dwells. For one-dwell,

Last Dwell Unchosen, First Dwell Left trials, there was a significantly greater proportion of Easy trials than Hard trials (*M* _*Hard−Easy*_ = -0.00921, *p* = .00421) and the opposite pattern for four or more dwell trials (*M* _*Hard−Easy*_ = 0.0138, *p <* .001). Taken together we see that Hard trials generally shift toward having more dwells (three or more) than Easy trials (one or two).

Shifting to the follow up analysis of the more infrequent trials where the First Dwell was to the Right, the only significant effects came from the main effects of Last Dwell Option and Dwell Count (Last Dwell Option: *F* (1,96) = 17.502, *p <* .001; Dwell Count: *F* (2.06, 197.43) = 24.566, *p <* .001). All the pairwise comparisons were significant, with a greater proportion of First Dwell Right and Last Dwell Chosen trials than First Dwell Right and Last Dwell Unchosen trials (*M* _*Chosen−Unchosen*_ = 0.00791, *p <* .001), and, when the First Dwell started on the Right, a pattern of greater proportions of two-dwells than one-dwell than three-dwells than four or more dwells (all *p*’s ≤.01689).

Figure 1C highlights the dominant gaze patterns evident during the Photo Preference task. Most often, participants looked at each option at least once, almost always starting on the left and ending on whichever option they chose. Decision difficulty inflated the number of dwells, with participants more likely to make more dwells if the decision was harder.

### 2.3 Gaze Dynamics: Driven early by stereotyped information gathering, affected later by decision difficulty

We used our proportion of trials analyses to guide an in depth exploration of gaze dynamics for the most commonly occurring gaze patterns in each task. This ensured adequate statistical power and allowed us to fully represent the dramatic ways gaze patterns differed across tasks. The analyses presented in the current section focused on the temporal aspects of gaze patterns and specifically examined how these dynamics varied with decision difficulty. We operationalized the gaze dynamics into timing metrics to describe the onset and duration of a dwell, as well as the time from the end of the dwell to the end of the trial (where the trial ended upon the cursor entering the chosen option). In cases where the most frequent gaze pattern involved more than 1 dwell, we also measured the onset and duration of the second or last dwell, as well as the time between dwell events. Across the three tasks, we analyzed 21 gaze dynamics metrics. We conducted paired t-tests comparing the Hard and Easy Difficulty conditions for each metric. To account for the potential impact of multiple comparisons, we adjusted the significance level to *p* = .05/21 = .00238. We fully report the results of these twenty-one gaze metrics in Table 2.

**Table 2:** Pairwise comparisons between hard and easy trials for all gaze metrics. Note. Gaze dynamics were tested at an alpha level of .05/21 = .0023809. *p *<* .0023809; **p *<* .0001; ***p *<* .00001

| Measure | *M* |  | Hard-Easy |  |  |  |  |
| --- | --- | --- | --- | --- | --- | --- | --- |
|  | *Decision Difficulty* |  | *M* | *SE* | df | *t* | *p* |
|  | Hard | Easy |  |  |  |  |  |
| Sentence Verification |  |  |  |  |  |  |  |
| 1 Dwell on Chosen |  |  |  |  |  |  |  |
| First Dwell Onset (ms) | 1857 | 1149 | 708 | 41.5 | 88 | 17.0 | *** |
| First Dwell Duration (ms) | 227 | 214 | 13 | 11.8 | 88 | 1.1 | n.s. |
| End Dwell to Response (ms) | 89 | 42 | 46 | 14.5 | 88 | 3.2 | * |
| Numeric-Size Congruity |  |  |  |  |  |  |  |
| 1 Dwell on Chosen |  |  |  |  |  |  |  |
| First Dwell Onset (ms) | 448 | 429 | 19 | 9.0 | 89 | 2.1 | n.s. |
| First Dwell Duration (ms) | 302 | 267 | 35 | 8.2 | 89 | 4.3 | ** |
| End Dwell to Response (ms) | 74 | 83 | -9 | 6.5 | 89 | -1.4 | n.s. |
| 2 Dwells with the Last Dwell on Chosen |  |  |  |  |  |  |  |
| First Dwell Onset (ms) | 263 | 263 | -0.5 | 8.3 | 87 | -0.1 | n.s. |
| First Dwell Duration (ms) | 207 | 217 | -10 | 4.9 | 87 | -2.0 | n.s. |
| Transition Duration (ms) | 150 | 140 | 10 | 6.4 | 87 | 1.6 | n.s. |
| Last Dwell Duration (ms) | 248 | 211 | 37 | 8.9 | 87 | 4.2 | ** |
| End Dwell to Response (ms) | 36 | 29 | 7 | 3.3 | 87 | 2.0 | n.s. |
| Photo Preference |  |  |  |  |  |  |  |
| 2 Dwells with the First Dwell on Left, and the Last Dwell on the Right Chosen |  |  |  |  |  |  |  |
| First Dwell Onset (ms) | 282 | 272 | 10 | 17.7 | 79 | 0.6 | n.s. |
| First Dwell Duration (ms) | 524 | 494 | 30 | 48.6 | 79 | 0.6 | n.s. |
| Transition Duration (ms) | 156 | 131 | 25 | 10.7 | 79 | 2.4 | n.s. |
| Last Dwell Duration (ms) | 719 | 514 | 205 | 35.6 | 79 | 5.7 | *** |
| End Dwell to Response (ms) | 51 | 39 | 12 | 8.9 | 79 | 1.3 | n.s. |
| 2 or More Dwells with the Last Dwell on Chosen |  |  |  |  |  |  |  |
| First Dwell Onset (ms) | 295 | 282 | 13 | 9.5 | 94 | 1.4 | n.s. |
| First Dwell Duration (ms) | 454 | 477 | -23 | 13.2 | 94 | -1.8 | n.s. |
| Last Dwell Onset (ms) | 1482 | 1261 | 221 | 31.2 | 94 | 7.1 | *** |
| Last Dwell Duration (ms) | 521 | 407 | 114 | 19.1 | 94 | 6.0 | *** |
| End Dwell to Response (ms) | 117 | 59 | 57 | 14.1 | 94 | 4.1 | ** |

#### 2.3.1 Sentence Verification Task

The most common gaze behaviour observed during the Sentence Verification task was either no dwell on any choice option or a single dwell on the chosen option. As mentioned earlier, we did not investigate gaze dynamics for the no dwell on option trials. To examine the gaze behaviour on single dwell trials, we analyzed three metrics: First Dwell Onset, First Dwell Duration, and End Dwell to Response (see Figure 2A). In the Sentence Verification task, irrespective of decision difficulty, there was no difference in the duration of the dwell on the chosen option. However, for both First Dwell Onset, and the time from the end of the dwell up to the response (End Dwell to Response), Hard decisions elicited significantly longer latencies (see Table 2). While we know that Response Times are longer for hard decisions (see above), these gaze dynamics suggest that the additional time is not simply due to an elongated timeline for all components of a Hard decision. Rather, in this particular task, where decision-relevant information resides outside of the choice options, more time is spent reading the sentence for Hard decisions before the gaze moves to the chosen options (approximately 700 ms). But, the chosen option itself does not require extensive viewing as the ‘True’ and ‘False’ options maintain a consistent position throughout the task. Hence this dwell time does not differ between Hard and Easy trials. However, after the gaze leaves the chosen option, a harder decision requires more time to complete compared to an easier decision. This suggests that there is further cognitive processing involved in resolving the decision before the mouse cursor lands in the chosen option. We discuss this finding in light of our parallel mouse-tracking study (Ouellette Zuk et al., 2023) in the Discussion.

#### 2.3.2 Numeric-Size Congruity Task

For Numeric-Size Congruity we analyzed the two most prevalent gaze behaviours: a single dwell on the chosen option, and two dwells with the last dwell on the chosen option. Interestingly, in contrast to Sentence Verification, the effect of difficulty on gaze pattern timing in single dwell trials was entirely different. Figure 2B illustrates that the only significant difference between Hard and Easy trials was found in the dwell duration of the single fixation, with Hard trials eliciting longer dwells than Easy trials (see Table 2). In this task, since the decision-relevant information is located at the choice option, the effects of decision difficulty are primarily expressed in the time required to visually acquire this information.

Further insights about the decision-making process are gained when exploring the two-dwell pattern with the last dwell on the chosen option. In these trials, we segmented the pattern into 5 constituent metrics: time to first dwell, first dwell duration, time between dwells, second dwell duration and second dwell offset to response. Remarkably, only the duration of the second (and last) dwell showed a significant effect of decision difficulty (see Table 2). This indicates that in two-dwell trials, both Hard and Easy trials exhibit the same initial dwell gaze pattern up until the second, and chosen, choice option is viewed. Only at this point, does the gaze tend to dwell longer on average for Hard decisions, suggesting that additional time is needed to process and integrate the decision-related information obtained from both options.

#### 2.3.3 Photo Preference Task

To capture the unique gaze behaviours specific to Photo Preference, we examined two distinct patterns: the highly stereotypic gaze pattern of two dwells with the first dwell on the left and the last dwell on the chosen (and right) option, as well as a broader set of patterns involving more than one dwell where the last dwell was on the chosen option. Similar to the Numeric-Size Congruity task, we analyzed five metrics for two-dwell, first-dwell-left, last-dwell-chosen trials. Once again, we found significant effects of difficulty only in the second (and last) dwell, with Hard trials resulting in dwells significantly longer than Easy trials (see Table 2). Figure 2C illustrates this pronounced difference, and we later discuss how the form of this decision information (colorful photos) and the type of decision required (preference) likely contribute to this finding. Lastly, we also examined decision difficulty in Photo Preference across all gaze patterns involving more than one dwell (trials with two, three or four or more dwells) where the last dwell fell on the chosen option. Here, we still analyzed five metrics, but with adjustments: metrics were anchored to the ‘last’ dwell (whether it was the second, third, fourth or more), and the measure of time between the two dwells was replaced with the onset time of the last dwell relative to the start of the trial. This is depicted in Figure 2C. In these analyses, we observed significant differences between Hard and Easy trials in the duration of the last dwell, the onset of the last dwell, and the offset of the last dwell to the response (see Table 2). Hard trials consistently exhibited longer latencies in all three metrics. As these tests were performed on aggregated data from two, three, four, or more dwells, the onset of the last dwell aligns with the results from the proportion of trials, where Hard Photo Preference trials had a greater proportion of trials with more dwells compared to Easy trials. Once again, the duration of the last dwell provided evidence that it was not exclusively the ‘second’ dwell but rather the ‘last’ dwell affected by decision difficulty. Further, the impact of decision difficulty extended beyond the last gaze, as Hard decisions appeared to require additional time for resolution, consistent with the longer movement times in Ouellette Zuk et al.’s study (2023). Perhaps most interestingly, just like the Size-Congruity task, the timing of the first dwell was not impacted by decision difficulty. That is, in terms of both onset and duration, the first dwell is highly stereotyped, suggesting that decision competition doesn’t really begin until all decision information has been sampled.

## 3 Discussion

In this study we collected data from a remote cohort of participants performing three different binary choice tasks while recording gaze behaviour via webcam eye-tracking. This paper serves as a companion to Ouellette Zuk et al. (2023) which details how decision-dynamics play out for the same three tasks when measured by mouse-tracking. Both papers rely on previously published works [10, 12, 26] that identified trials with hard or easy decisions. Briefly the three tasks were: A Sentence Verification task where you determined whether a statement was true or false (difficulty was manipulated through sentence negation);

A Numeric-Size Congruity task where you determined which of two digits was numerically larger (difficulty was manipulated through the congruence of physical and numerical size); and a Photo-Preference task where you determined which of two photos you preferred (difficulty was manipulated by the pleasantness-similarity between the two photos). In general, hard choices take longer to resolve than easy choices, a finding we report in the companion paper and replicate here in our analysis of Response Times (see Figure 1). However, response time is a coarse measure that cannot reveal any of the underlying decision processes. Similarly, though it is able to fill in the gap about the effects of decision-difficulty on decision processes *after* a movement is initiated, mouse-tracking (e.g., Ouellette Zuk et al., 2023) is blind to decision processes arising *prior* to movement onset. Thus, the primary objective of this study was to examine how decision difficulty manifests in gaze dynamics - a measure capable of indexing decision processes from the moment of stimulus onset.

In order to investigate gaze dynamics it was first necessary to more broadly categorize gaze patterns - the series of looks (dwells) the eyes made on certain targets relevant to a decision. For example, it would be meaningless to examine the dynamics of a second-dwell for a task where this rarely occurred. We first examined trial characteristics including number of dwells, the side of space of the first dwell and whether the last dwell was toward the chosen target, calculating the proportion of trials observed for each gaze pattern and whether they occurred in hard or easy trials. This analysis of common patterns first revealed a task-general difficulty effect whereby more dwells were observed on harder trials. Second, and more importantly, it also highlighted distinct and unique gaze patterns observed *between* tasks. While the first paper in this series (Ouellette Zuk et al., 2023) demonstrated the consistency of decision difficulty effects between tasks, here the proportion of trials analysis exposed a relationship between gaze behaviour and the spatial distribution of decision information within a task.

In the Sentence Verification task all of the decision information is contained in a statement at the top-middle of the screen with no unique information contained at the left or right choice options (the “True” and “False” labels at these locations remained constant). As such, the dominant gaze behaviour in this task contained either no looks toward the choice options or a single look toward the option that was selected. On the trials with one dwell on the chosen option, we found that the duration of that dwell remained the same for both hard and easy trials. Instead, the effects of decision difficulty emerged in the time it took for the dwell to start, and the time from the end of the dwell to the response. In this task we can infer that the gaze was focused on the sentence, and as the sentence contained all of the decision information, the differentiation between hard and easy trials emerged prior to any look towards the response options. In the case where a subsequent look to the chosen target occurs, the constant dwell duration seen regardless of decision difficulty suggests that this gaze might only serve the difficulty-independent process of spatially guiding the mouse response. However, after the gaze leaves the chosen option, difficulty effects re-emerge, suggesting that additional cognitive processing may be required beyond the choice-option dwell to finalize the decision.

In the Numeric-Size Congruity task, unlike Sentence Verification, the information necessary to make a decision is contained at the choice options. However, this task has the interesting property that sometimes a single fixation toward a choice option is sufficient to make a decision (e.g., if your eyes dwell on the digit “1” or “9” you can definitively know it is the lower or higher numeric value respectively) while other times a single dwell is insufficient (e.g., if you eyes dwell on the digit “2” or “8” fixating the other target is necessary to make a definitive decision). Given this, it is logical that the dominant gaze patterns observed in this task are divided into trials with only a single dwell and trials with two dwells. When examining the single dwell on chosen trials of the Numeric-Size Congruity task, difficulty effects manifest in a manner wholly opposite to that of the Sentence Verification task. Specifically, when participants focus only on the chosen option, they spend significantly more time dwelling on it during hard trials compared to easy trials. This prolonged dwell on the incongruent yet correct choice option is the only metric in the Numeric-Size Congruity task where difficulty effects become evident. Neither the onset of the first dwell nor the offset to response show any changes with difficulty. This begins to fill in the picture of how eye gaze functions with respect to decision difficulty - at the moment when sufficient information about the decision has been acquired via eye gaze then, and only then, does difficulty begin to differentially affect the decision timeline.

This hypothesis of gaze distribution being tied to information acquisition receives additional support from the two-dwell trials in the Numeric-Size Congruity task. Here we observe an intriguing pattern whereby the gaze dynamics of hard and easy trials appear identical all the way until the second dwell, where then, and only then, do hard trials exhibit a significantly longer duration for the last dwell compared to easy trials. Following this extended second dwell, the time to the response is comparable for both difficulty conditions. Thus, all the effects of decision difficulty for this set of trials are reflected exclusively in the last dwell, where determining the correctness of a numerically large but physically small (i.e., incongruent) choice option requires more dwell time compared to a numerically large and physically large (i.e., congruent) choice option. Perhaps most importantly, the first dwell, which is predominantly directed towards the unchosen option, does not take longer in the hard condition. It is only when participants view both choice options, and therefore have acquired all the necessary decision information, that difficulty effects emerge.

Finally in the Photo Preference task, decision information is necessarily and evenly distributed between both choice targets. That is, you can’t make a determination of comparative preference between two photos without having your eyes dwell on both of them. Accordingly, when examining the most common gaze patterns in the Photo Preference task, they all involve two or more dwells with the single most common pattern being a look to the left, unchosen target followed by a look toward the right, chosen target. Focusing on this specific two-dwell pattern, the results follow the structure outlined in the Numeric-Size Congruity task. That is, the difficulty effect emerges exclusively in the last dwell duration of these Photo Preference trials. Like before, a hard trial does not exhibit signs of being difficult until the second, chosen option is viewed.

These results are confirmed in our broader investigation of Photo Preference trials with two or more dwells (e.g., trials with two, three, four or more dwells). Again, we consistently observed difficulty effects in the latter half of the trial, with hard trials showing a longer duration for the last dwell, as well as later onsets of the last dwell and longer durations between the last dwell and the response. This broader result shows the consistency of the difficulty effect on the duration of the last dwell, while the prolonged onset of the last dwell on hard trials likely results from averaging trials with a higher dwell count. Of note, the time from the offset of the last dwell to the response should not be conflated with the number of dwells in the same way. Instead, it suggests that this broader set of trials, with more dwells in harder trials, may more clearly capture decisions with lingering uncertainty. In other words, if harder trials take longer and more dwells cost more time, this set of trials may reflect the more challenging choices where difficult decisions are still being resolved all the way until the final choice occurs.

Taken together, the analysis of gaze behaviour across these three binary choice tasks sheds light on the dynamics of decision making. It appears there are at least two processes at play - the gathering of decision information and the resolution of the decision. While these two aspects of the decision can likely proceed in parallel, our data suggest that information gathering is the predominant driver of gaze early in trials and proceeds largely without impact from the specific demands of decision difficulty until all the relevant information has been at least partially sampled. Moreover, information gathering appears to be highly stereotyped for a given task. This is most evident in the Photo Preference trials - when information was evenly distributed across two locations, the vast majority of dwells were directed first to the left, and then to the right. This aligns with gaze-focused decision models trained on behavioral data from similar binary choice preference tasks [28, 42, 5]. These models assume a left-first gaze, highlighting the persistent influence of ingrained eye movements used in reading left to right (at least in the English-speaking population tested). Additional evidence for the stereotyped nature of information gathering is seen in the first dwell dynamics on trials with more than one dwell (Numeric-Size Congruity and Photo Preference). Here, the time to first dwell and first dwell durations are not impacted by decision difficulty.

This suggests that there is a level of dynamics involved in decision making that most models don’t capture. That is, most decision-models simulate a dynamic decision using a series of static parameters - for example the rate at which evidence for an option accumulates and the bound to which it must accumulate to in order for it to be chosen. However, here we show that the parameters themselves are likely changing throughout a decision. Specifically, we argue that the value of a given choice option necessarily fluctuates as the decision shifts from information gathering to decision resolution. Consider the Photo Preference task. Initially a target’s value is dictated by its ability to deliver new information (for a review of this idea, see [20]), separate from its content (e.g., pleasantness). Thus, both targets are equally valuable and the eyes adopt a stereotyped left-to-right pattern of information sampling. But then, target-value shifts to being defined by the details of the image. Now, the pleasantness of each option, and critically the relative pleasantness between options, dictates the resolution of the decision. This ability to shift decision parameters based on the current task context can explain both of our major gaze dynamic findings: initial dwells are stereotypical, driven by information gathering and not affected by decision difficulty, while the last dwells are affected by decision difficulty since value during decision resolution is determined by the relative difference in task-relevant content between targets.

Understood this way, it is clear that multiple facets of a target determine its value: if it contains task-relevant information, if it has been looked at, if it is an image or text, if it is small or large, if it is easily identified or not. Equally clear is that which facets are of value changes over time. Across our two studies we can measure the value transition from information gathering to decision resolution - a specific example of the more general pattern of explore versus exploit behaviors [8]. Since a key aspect of exploration is the physical location of targets, eye-tracking is particularly well suited to measure these kinds of information gathering behaviours. Then, as a decision shifts to resolution, which typically demands a motor response, mouse-tracking becomes a sensitive tool for watching the later stages of competition play out across time and space. Together, this highlights the complementary approaches taken across our two companion papers: gaze behaviour is acutely sensitive to task differences, especially early in trials and with respect to the spatial distribution of decision-relevant information, while mouse tracking is more sensitive to the decision difficulty effects that appear once all relevant information has been sampled.

We believe a cross-paper comparison of difficulty effects between tasks offers some initial support for this idea. Arguably, the gaze dynamics during Sentence Verification are the least informative - many trials have no dwells at the site of a choice and those that do, don’t have dwell durations that differentiate between hard and easy trials. In complete contrast, as we report in our companion paper (Ouellette Zuk et al., 2023), during-movement measures of movement time and trajectory strongly differentiate between hard and easy trials for the same task. On the other end of the spectrum, the current analysis of gaze patterns in the Photo Preference task offers rich information about the contemplation of decision options, including many trials with multiple dwells. But, this same task in our companion paper (Ouellette Zuk et al., 2023) shows that during-movement measures had, relative to the other tasks, the least sensitivity to decision-difficulty. Not only does this show why collecting gaze and movement data is important, it also has theoretical implications. We previously made the distinction between gathering and resolving decision information - related processes that can sometimes proceed in parallel. We speculate that tasks like Photo Preference which require longer times spent gathering information are thereby also granted extra time to start resolving a decision prior to movement onset. As a result, movement related measures in these kinds of tasks show less sensitivity as more of the decision has been completed prior to the initiation of a response.

Aside from their impressive combined ability to cover the full range of a decision - from stimulus onset to response completion - there is another, methodological link between our two companion papers: their use of remote data collection. Not only did it vastly increase the sample size of our studies (more than 400 data sets initially collected across the two papers) it also made the study accessible to participants who may not typically participate in academic research (see supplementary material in Ouellette Zuk et al., 2023). Maybe most importantly, it also allowed us to test participants in more ecologically-valid environments. That is, we can collect data from people using their own devices from the comfort of their own homes without introducing an artificial, isolated, and highly-controlled laboratory setting that likely limits our ability to capture realistic and natural human behaviour [25, 41].

## 4 Limitations and Conclusion

Together, this paper series demonstrates the power of remote, online methods as tools for deeply understanding the complete, dynamic and continuous decision process, rewinding the decision from the typically-collected final choice response, all the way back to the first glance. Combined, our studies reveal an incredibly robust set of findings, on smartphones, tablets and computers with mouse-tracking or eye-tracking across three unique decision tasks we can measure and decompose the effects of decision difficulty with precision.

Of course, full remote data collection is not without its limitations, some which are particularly evident in the current study. Most notably, there are known spatial and temporal inaccuracies when using webcam eye-tracking. As an example, we did not feel we had sufficient spatial accuracy to properly analyze nuanced reading behaviour during the initial dwells to the statement in the Sentence Verification task. This limitation also led to an unforeseen and unfortunate outcome - due to the sampling rate slowdown caused by prioritizing the collection of webcam eye-tracking, our mouse-tracking data in the current study was not sufficiently sampled to perform a confirmatory analysis of the effects reported in our companion paper (Ouellette Zuk et al., 2023). Finally, at present, webcam eye-tracking is generally restricted to participants using a computer with a webcam - it hasn’t yet been widely used with sufficient accuracy on mobile devices in decision science research.

These limitations, however, are the type which seem likely to be overcome soon. Improved and more efficient gaze detection driven by ever-improving machine learning models, continued advancements in consumer hardware, and the use of mobile device cameras for eye-tracking research (e.g., [34]) are all on the imminent horizon [24]. And in that future, we envision using gaze and movement tracking will be paramount to understanding participant behaviour in and out of the lab. Importantly, as we advocate for moving into unrestricted domains - where the nature of the task and the decisions that people make are not controlled by an experimenter - we recognize the need for and strength of this combined metrics approach. Including both gaze and movement analysis allows you to understand both where the most relevant decision information is located via eye-tracking, and via motion-tracking, how difficult it is to adjudicate between that information to arrive at and perform the final movement required to enact a choice.

## 5 Methods

### 5.1 Participants

100 adults provided their informed consent to participate in the experiment, and completed the study in full. Of the 100 participants, 36 self-identified as female, 66 as male, and one participant preferred to not disclose their gender. The average participant age was 25.47 years old (+/- 4.28). Participants were recruited using Prolific (www.prolific.co), an online crowdsourcing platform, where we followed Ouellette Zuk et al.’s (2023) participant restrictions on age (18 to 35 years old), and prior approval rating on the platform (95-100%). We paid participants for their time (6 GBP per hour, ∼$10 CAD per hour). All experimental proceedings were approved by the University of Alberta’s Research Ethics Board (Pro00087329) and were performed in accordance with relevant guidelines and regulations.

### 5.2 Materials

All participant data for the study was collected through the use of Labvanced [13], an online, browser-based Javascript experimentation platform. We designed our study in a 800 × 450 pixel coordinate frame in Labvanced (see Figure 3B), where it would automatically scale to the size of the participant’s screen. We used Labvanced’s built-in webcam eye-tracking (Labvanced v2 High Sampling Mode eye-tracking [13]). To ensure high-quality data collection, certain minimum requirements were imposed. Participants were required to use either a laptop (n = 75) or desktop (n = 25) computer with a mouse. The operating systems supported were Mac (n = 15), Windows (n = 84) or Linux (n = 1), along with the Chrome browser. Furthermore, participants were required to have a webcam with a minimum resolution of 1280 × 720 pixels, a landscape-oriented screen with a minimum resolution of 600 × 600 pixels (*Mode* = 864 × 1536 px), and a computer system capable of collecting at least 10 samples per second of the head’s position for optimal eye-tracking precision (*M* = 15.65 Hz, *SD* = 5.69 Hz). This system threshold was often met if participants had a graphics card and had freed up system resources prior to starting the study (i.e. closing any other programs running on their computer).

### 5.3 Task & Procedure

Broadly, the tasks and procedures followed by Ouellette Zuk et al. (2023) were repeated here but with the addition of webcam eye-tracking, and with only computers being included (Ouellette Zuk et al also tested tablets and smartphones). Like Ouellette Zuk et al. (2023), we asked participants to complete three distinct tasks that all required decision-making in a binary choice paradigm where choices were made with mouse movements. These tasks were Numeric-Size Congruity, Sentence Verification, and Photo Preference (with examples shown in Figure 1).

Participants recruited through Prolific (www.prolific.co) were given access to a detailed description of the study. This description included an approximate duration of the study (1 hour), information regarding the necessary hardware, and instructions on how to avoid any potential technical difficulties (complete Prolific description available in the Supplementary Materials). Upon clicking the study link that accompanied the study’s description, participants were directed to a full-screen Labvanced browser window and prompted to grant permission for their webcam device. In the event that participants did not meet the minimum requirements, they would immediately receive an error or warning message. Assuming no issues arose, participants would begin by providing their informed consent to participate, after which they would proceed to answer a brief survey pertaining to their demographic information and the hardware they were using.

Before the primary experimental tasks, participants were given information to encourage successful webcam eye-tracking data collection. Instructions about optimal lighting conditions, the re-calibration process, and the virtual chinrest feature were provided to participants. Then, participants underwent Labvanced’s 5-minute eye-tracking calibration procedure. It was required that participants redo the calibration if the predicted gaze error surpassed 7% of the screen’s dimensions. After completing calibration, participants began the main experiment.

Figure 3 illustrates the experimental procedure and trial progression. Three tasks were performed (counterbalanced in their presentation order), with simple task instructions preceding each task. 84 trials were completed per task, with task stimuli presented in a randomized order within each task. Every 5 trials, a brief seven-point eye-tracking recalibration procedure was performed to adaptively correct any drift errors in the gaze prediction algorithm over the course of the experiment. At any point during an experimental trial, the virtual chinrest feature would pause the trial if a participant’s excessive head movement affected the quality of the gaze prediction (*M* = 6.59 trials, *SD* = 7.22 trials).

An experimental trial began with a green circular start button labeled “Touch here” at the bottom center of the screen. Participants had to move their mouse cursor to the button to initiate the trial. This prompted the appearance of a three-second countdown at the center of the screen. If the mouse cursor was removed from the circular button, the countdown paused until the cursor’s return. In the Numeric Size-Congruity and Photo Preference tasks, a task-specific question appeared at the top center of the screen during the countdown (see Figure 1). Once the countdown ended, two choice boxes appeared at the upper-left and upper-right corners of the screen, presenting trial-specific options. In the Sentence Verification task, two choice options appeared alongside the countdown and displayed a statement at the top center of the screen after the countdown completed. Participants could immediately select their choice option by moving their mouse cursor inside the respective choice area. Once a choice was made, the selected box was highlighted while the other choice option and start button disappeared. A “Next” button then appeared at the center of the screen for participants to click to proceed to the next trial at their own pace.

All three tasks required binary choice decisions in Hard and Easy conditions. In the Numeric-Size Congruity task, participants were presented with pairs of digits and asked to determine which digit had a higher numeric value. The pairs of digits varied in congruence, where some pairs were congruent in both numeric and physical size (representing Easy trials with low decision difficulty, e.g., 2 vs. 8), while others were incongruent in numeric and physical size (representing Hard trials with high decision difficulty, e.g., 2 vs. 8). In the Sentence Verification task, participants were tasked with verifying the truthfulness of statements. From previous work [10] it has been shown that statements that are true show large decision difficulty effects based on whether they are non-negated (representing Easy trials with low decision difficulty, e.g., ‘Cars have tires’) or negated (representing Hard trials with high decision difficulty, e.g., ‘Cars do not have wings’). In the Photo Preference task, participants were presented with pairs of photos that differed in valence (from the International Affective Picture System stimulus set [31], as in [26]). They were asked to then choose which photo they preferred. The pairs of photos varied in their dissimilarity of valence, with some pairs being dissimilar (representing Easy trials with low decision difficulty, e.g. High vs. Low pleasantness) and others being similar (representing Hard trials with high decision difficulty, e.g. High vs. High pleasantness). These tasks were designed to cover a wide range of decision domains, including objective perceptual judgments (such as discriminating between digits), semi-subjective conceptual judgments (such as evaluating the truth value of statements), and subjective preference judgments (such as expressing a preference for specific photographs). Additionally, the tasks intentionally differed in terms of stimulus characteristics and the cognitive processing requirements involved.

The entire experimental procedure, as a Labvanced study, can be accessed via the link in Supplementary Materials.

### 5.4 Data Processing

The uncontrolled nature of online, remote data collection, including the use of webcam eye-tracking, presented some data quality challenges that required thoughtful treatments. Gaze and cursor timeseries data were collected in a way that maximized the number of data samples processed for each participant, with priority given to the collection of gaze samples. All gaze and cursor data were then upsampled (linearly interpolated) to a common sampling rate of 60 Hz. The gaze prediction algorithm was refined (i.e., recalibrated) every 5 trials using Labvanced’s adaptive drift correction method [13], therefore gaze data quality varied over time. To minimize data rejection, we assessed the quality of each participant’s gaze data within each task independently (as opposed to rejecting an entire dataset for poor gaze data for a subset of trials). Using custom MATLAB scripts and our Gaze and Movement Analysis software, our initial data cleaning approach aimed to assess whether the gaze data showed reasonable patterns or whether it contained noisy, spurious gaze prediction errors (see [4] for a similar approach). Using more exaggerated but still mutually-exclusive boundaries (see Figure 3B - AOI Dwell Boundaries), we determined whether the gaze fell inside at least one of the task-critical areas during the decision period. For the Numeric-Size Congruity and Photo Preference tasks, we considered the gaze data of reasonable quality if the gaze fell within the left or right choice options for at least 100 milliseconds continuously. For the Sentence Verification task, we instead used at least one look (minimum 100 ms) to the sentence area as a proxy of reasonable gaze data as it was common for participants to not look at the choice options. In each task, if less than half of the trials showed reasonable gaze data (i.e., one dwell on a task-relevant AOI), we removed the full task from the participant’s dataset for analysis. Based on this criteria, from 100 participants, 3 subjects’ Photo Preference task data were removed, 9 subjects’ Numeric-Size Congruity task data were removed, and 3 subjects’ Sentence Verification task data were removed. Of the remaining datasets, any individual trials that failed to show reasonable gaze data (per the same criteria) were removed leaving the Photo Preference, Numeric-Size Congruity, and Sentence Verification tasks with 82.79 (+/-2.96), 73.41 (+/-9.58) and 81.70 (+/-5.79) of 84 trials respectively.

Then, to assess non-gaze-related quality of data, we employed similar rejection criteria to Ouellette Zuk et al. (2023). Within each task, we removed any trials where the response time was less than 100 milliseconds or greater than 3 standard deviations above the subject’s mean response time in that task. We also removed any trials where a pause occurred from the virtual chinrest feature, or any trials where the response time was not computable (from participant error or occasional data recording issues). Further, any incorrect trials were removed from the Sentence Verification and Numeric-Size Congruity tasks (where accuracy could be assessed objectively). Following these additional trial rejections, we again removed entire task data from any participant with less than 42 or their original 84 task trials, which resulted in one additional Numeric-Size Congruity task removal. From 100 participants, the final task datasets included in analysis were: Photo Preference: n = 97, 78.30 (+/- 5.18) trials, Numeric-Size Congruity: n=90, 70.87 (+/-9.16) trials, Sentence Verification: n=97, 72.09 (+/-8.23) trials.

Unlike our companion paper (Ouellette Zuk et al. 2023) where cursor-tracking provided the key dependent measures, high-quality gaze data from webcam eye-tracking was the primary goal of the current study. While we did record mouse trajectory data alongside our gaze data, we did not have sufficient data quality to reliably metricize the mouse trajectory data in the same way as Ouellette Zuk et al. (2023). With recording priority given to the gaze data, and cursor movements generally being very quick, we found that there were many instances that the number of data samples from the cursor were insufficient. We discuss this unintended consequence in the Discussion, and encourage readers to engage with our companion paper (Ouellette Zuk et al., 2023), which provides a thorough analysis of high-quality cursor and touchscreen trajectories from an original sample of more than 300 other participants.

### 5.5 Dependent Measures

Our analysis strategy revolved around three sequential steps. First, to confirm and replicate that the main decision-difficulty effects elicited by these tasks were present in the current study, for every trial we recorded *Response Time (ms)*: the time from the choice options being presented to the moment the cursor was detected as entering within the bounds of a choice option.

Second, to broadly characterize and analyze the dominant gaze patterns within each task, we calculated *Proportion of Trials (%)*: a value of 0 or 1 for each trial that represented if that trial shared certain characteristics (e.g. was it Hard or Easy). These counts were then aggregated such that proportion of trials for a given characteristic was always calculated within each task per participant, where the trials fitting the characteristic were counted and then divided by the total number of trials included in the analysis (specifically, the number of Hard plus Easy trials).

Third, we used the results from the proportion analysis to guide our examination of the gaze dynamics of the most frequent gaze patterns within each task. These gaze dynamics were centered on describing the timing of specific dwell patterns - when a dwell started, how long it lasted, and when it ended relative to the response. For patterns with more than one dwell, we wanted to capture this information about both the first and last dwell. Using the first and last (as opposed to strictly second) dwell afforded us flexibility in describing two-dwell patterns, but also three, four or more dwell patterns. In all three tasks, for every gaze pattern observed (where every gaze pattern necessitated there being at least one dwell), the following measures were collected:

*First Dwell Onset (ms)*: the time from the choice options being revealed to the start of the first dwell within a choice option.

*First Dwell Duration (ms)*: the length of time the dwell stayed at the first choice option viewed.

*End Dwell to Response (ms)*: the time from the end of the final dwell to the response, as determined by the mouse cursor’s entry into the choice option.

When a frequent gaze pattern included more than one fixation (only the Numeric-Size Congruity and Photo Preference tasks), the last dwell’s duration was also collected:

*Last Dwell Duration (ms)*: the length of time the dwell stayed at the last choice option viewed preceding a response.

To capture the time of the last dwell’s onset, we used two measures, dependent on the number of dwells in the gaze pattern being explored. When looking at trials with exactly two dwells, we use Transition Duration as a measure, but when looking at a collection of trials with two or more dwells (Photo Preference only), we use Last Dwell Onset:

*Transition Duration (ms)*: the time between the first and second (i.e. last, in these cases) dwells, measured from the offset of the first dwell to the onset of the second dwell.

*Last Dwell Onset (ms)*: the time from the choice options being revealed to the start of the last (i.e. second, third, fourth or more) dwell within a choice option.

### 5.6 Statistical Procedure

Each dependent measure was analyzed within each task using Jamovi (Version 2.2.5; an open-source statistical software). Response times and gaze dynamic metrics for each task were analyzed using paired t-tests of each participant’s Hard and Easy trial means on that given task. To correct for multiple comparisons, the three response time tests were performed with an alpha of .05/3 = .01667, and the twenty-one gaze dynamic metrics were tested with an alpha of .05/21 = .00238. The proportion of trials measures were tested using Repeated Measures ANOVAs, where p-values were Greenhouse-Geisser-corrected for sphericity violations. The Sentence Verification task was tested with a 3 factor RMANOVA, and Numeric-Size Congruity and Photo Preference tasks were tested with 4 factor RMANOVAs. We followed the family-wise error correction procedure from Cramer et al. [9], where the threshold for significance becomes increasingly more conservative with every significant test result within a family of results. We treated all 3 omnibus RMANOVAs as a single family to determine the significance of the omnibus results for the proportion of trials measures. Follow-up RMANOVAs were then performed on the highest order interaction(s), testing each level of one factor against the other factors (see section 2 - Results). This interaction procedure was performed as necessary until a single-factor RMANOVA was reached, where the simple main effects of one factor could be tested at all levels of the other factor. Significant main effects were explored with all pairwise comparisons. The Cramer et al. [9] procedure was again employed for these follow-up RMANOVAs, where the family-wise error correction was performed within-task (i.e. each task’s follow-up tests became a family). We report our results in Tables 1 and 2. Table 1 presents the per task trial proportion results from the breakdown of each three-way interaction involving the factor of difficulty, and Table 2 fully reports the gaze metrics differences between the Hard and Easy trials for each of our 21 gaze-dynamic measures.

## Supporting information

Complete Supplementary Materials

## Acknowledgments

This study was funded by an NSERC Discovery Grant to CSC, and a MITACS Accelerate International award to JKB.

