## Supplementary material for "Continuous Measures of Decision-Difficulty Captured Remotely: II. Webcam eye-tracking reveals early decision processing": Complete Supplementary Materials

### Supplemental Materials 1 - Prolific Description

**Choose between 2 presented options -with webcam-based eye-tracking (*need high-perf. comp, MOUSE ONLY, graphics card*)**

By ualberta.ca

£6.50 • £6.00/hr | 65 mins | 30 places

Welcome to the experiment! This study should take approximately 65 minutes, and needs to be completed in one sitting. You will be required to stay **very still** during the entirety of the experiment for eye-tracking purposes.

***Some people may not have the internet speed, graphics card, and/or hard drive system performance that this study requires - many factors contribute to this.*** **If you've closed other internet tabs etc. and you still get an error that says the face/video processing is not working well enough, you will not get past the very first screen. You will be unable to participate and will need to RETURN THE STUDY. PLEASE DO NOT KEEP TRYING - it will not work.**

**A WIRED OR WIRELESS COMPUTER MOUSE IS ALSO REQUIRED. NO TRACKPADS/TOUCHPAD/TOUCHSCREEN/STYLUS.** Do not try - it will not work for the experiment (please do not waste your time trying). **Thank you!**

We are conducting an academic study about decision-making. To study this, we need participants to choose between two presented options on the computer while a webcam-based eye-tracking system determines where the participants eyes are looking. While you complete this decision task, your eye gaze position will be recorded. While the webcam is used to calculate your eye gaze position in numerical form, no webcam video data is recorded.

Warning: some images shown may contain potentially explicit or offensive content (e.g. blood, violence) that some viewers may find disturbing. Participant discretion is advised.

We appreciate your interest in participating! This scientific data will be used to help us develop a better understanding of the eye-gaze in decision-making.
Still interested? Here's the important details:

### **Very important - please read the following requirements carefully**

**To participate, make sure you:**
**-- Use Chrome as your browser**
**-- <u>Use a computer MOUSE (NOT A TRACKPAD OR LAPTOP TOUCHPAD ETC - WILL FAIL)</u>**
**-- Use only a laptop or desktop computer (tablets/phones will fail)**
**-- Can do a computer task without needing to wear glasses (contacts OK)**
**-- Have a connected webcam (either built-in or external device is fine)**
**-- Have your laptop or desktop computer on a stable table top (not on your lap), in a well-lit room - with light in front of you (not behind) so that your face isn't shadowed. Cannot be in a dark room with only light coming from the computer screen (eye-tracking system will fail)**
**-- Are aged 18 to 35**
**-- Have the ability to keep a consistent, still head position for the full duration of the study**
**-- Can keep the study full screen for the full study duration (approx. 60 mins)**
**-- Close all other internet tabs before starting the study (experiment will fail)**

The task will take approximately 60 minutes to complete. Complete the task in one sitting. You will not receive payment if you do not complete the task fully and in one sitting.

**DO NOT** exit from full screen mode during the experiment. If you are not using the Chrome browser, please swap browsers before beginning the study. The study will take 2-3 minutes to load at the beginning of the study.

**VERY IMPORTANT: Please close all tabs before beginning the study. You will receive an error if your background activity is impeding the performance requirements of the eye-tracking system and you'll need to return the study if the performance threshold is not met at the beginning of the study (you will get an error screen that tells you this). Some people may not have the internet speed or system performance that this study requires. If you've closed other internet tabs etc. and you still get an error that says the face/video processing is not working well enough, you will not get past the very first screen. You will be unable to participate and will need to return the study - do not keep trying. Thank you.**
-------
This research is carried out by researchers from the University of Alberta, in Edmonton, Alberta, Canada. The plan for this research has been reviewed for its adherence to ethical guidelines and approved by Research Ethics Board 2 at the University of Alberta (Pro00087329).

Devices you can use to take this study: Desktop
You will also need: Camera

### Supplemental Materials 2 - Labvanced Study Link

Experimental Task on Labvanced (for viewing purposes): https://www.labvanced.com/page/library/48965
